# Syntax-sensitive regions of Broca’s area and the posterior temporal lobe are differentially recruited by production and perception

**DOI:** 10.1101/2020.06.06.138131

**Authors:** William Matchin, Emily Wood

## Abstract

Neuroimaging studies of syntactic processing typically result in similar activation profiles in Broca’s area and the posterior temporal lobe (PTL). However, substantial functional dissociations between these regions have been demonstrated with respect to lesion-symptom mapping in aphasia. To account for this, Matchin & Hickok (2020) proposed that both regions play a role in syntactic processing, broadly construed, but attribute distinct functions to these regions with respect to production and comprehension. Here we report an fMRI study designed to test this hypothesis by contrasting the subvocal articulation and comprehension of structured jabberwocky phrases (syntactic), sequences of real words (lexical), and sequences of pseudowords (phonological). We defined two sets of language-selective regions of interest (ROIs) in individual subjects for Broca’s area and the PTL using the contrasts [syntactic > lexical] and [syntactic > phonological]. We found robust significant interactions of comprehension and production between these two regions at the syntactic level, for both sets of language-selective ROIs. This suggests a core difference in the function of these regions: language-selective subregions of Broca’s area play a role in syntax driven by the demands of production, whereas language-selective subregions of the PTL play a role in syntax driven by the demands of comprehension.

## Introduction

Although sentences appear as linear sequences of words, they are combined into hierarchical structures that determine their semantic interpretation (Chomsky 1957; Heim and Kratzer 1998). During online processing, syntactic mechanisms group words incrementally into these hierarchical structures (Crocker 1996; Schneider 1999; Lombardo and Sturt 2002; Sturt and Lombardo 2005). Neuroimaging studies of syntactic comprehension, such as the contrast of structured phrases or sentences to unstructured word lists, have revealed increased activation in a variety of brain regions (for a meta-analysis, see Zaccarella et al. 2017). These activations appear to be selective for higher-level linguistic computations, as these areas, when defined in individual subjects, do not show increased activation for a variety of non-linguistic tasks (Fedorenko et al. 2011). By contrast, there are spatially adjacent domain-general regions that appear to respond to a variety of non-linguistic tasks (Fedorenko, Duncan, et al. 2012; Fedorenko et al. 2013).

However, the comparison of noncanonical to canonical sentence structures, thought to tax syntactic processing resources (see Matchin and Rogalsky in press; Rogalsky and Hickok 2011 for discussion of this assumption), has primarily revealed activation in two areas: Broca’s area and the posterior temporal lobe (PTL) (for a meta-analysis, see Meyer and Friederici 2016). These two regions also selectively show jabberwocky structure effects, that is increased activation for structured sentences with content words replaced with pseudowords (greatly reducing conceptual-semantic content) relative to scrambled jabberwocky sentences (Pallier et al. 2011; Fedorenko, Nieto-Castañon, et al. 2012; Goucha and Friederici 2015; Matchin et al. 2017). Finally, they also show increased activation for verb phrases (with more complex syntactic organization) relative to lexically-matched noun phrases (with less complex syntactic organization), whereas other language-responsive regions do not exhibit this difference (Matchin, Liao, et al. 2019). This suggests that language-selective portions of Broca’s area and PTL support syntactic processing, broadly construed, whereas other regions more likely reflect semantic processes (Binder 2017; Matchin and Hickok 2020; Pylkkänen 2020).

However, neuroimaging studies have yet to ascertain a clear distinction of function between Broca’s area and the posterior temporal lobe. Individual neuroimaging studies of syntactic processing have sometimes reported isolated syntactic effects in Broca’s area without corresponding posterior temporal lobe effects (Stromswold et al. 1996; Caplan et al. 2000; Goucha and Friederici 2015; Zaccarella and Friederici 2015), but other studies reveal that both regions reliably exhibit these effects (Ben-Shachar et al. 2003, 2004; Bornkessel et al. 2005; Rogalsky et al. 2008; Obleser et al. 2011; Pallier et al. 2011; Fedorenko, Nieto-Castañon, et al. 2012; Matchin et al. 2017). Therefore, from the perspective of neuroimaging, it is unclear what the functional dissociation of these regions is with respect to syntax.

While the neuroimaging literature does not provide clear evidence of a functional distinction between Broca’s area and the PTL, lesion-symptom mapping analyses in aphasia have revealed distinct syntactic deficits following damage to these regions. Damage to PTL is associated with sentence comprehension and syntactic perception deficits, when confounding effects of working memory resources are accounted for (Dronkers et al. 2004; Wilson and Saygın 2004; Pillay et al. 2017; Rogalsky et al. 2018; Matchin and Hickok 2020). In addition, (Matchin et al. 2020) found that agrammatic production deficits (overall omission of functional elements and simplification of sentence structure) are associated with damage to Broca’s area but not PTL, whereas paragrammatic production deficits (grammatical errors with no overall omission/simplification) are associated with damage to PTL but not Broca’s area. These results led Matchin & Hickok (2020) to suggest that there is in fact a functional dissociation between these regions with respect to syntax, such that both areas are critically implicated in production, but only PTL is critically implicated in perception. However, while there are numerous neuroimaging studies of syntactic perception, few studies have attempted to isolate morpho-syntactic aspects of production (Haller et al. 2005; Del Prato and Pylkkänen 2014; Schönberger et al. 2014; Matchin and Hickok 2016) or directly comparing production and comprehension of syntax within the same study (Menenti et al. 2011, 2012; Segaert et al. 2012, 2013). This may account for the relatively limited neuroimaging evidence for distinctions in syntactic activation patterns between Broca’s area and the PTL with respect to production and perception of syntax.

In the present study, we decided to test the hypothesis offered in Matchin & Hickok (2020) by assessing syntactic perception and production in the brain in the same study. We used simple linguistic materials consisting of sequences of two-word jabberwocky structures (e.g., *this pand these clopes*), and a “perceive & rehearse” paradigm used in previous studies to localize both speech production and perception (Buchsbaum et al. 2001; Hickok et al. 2003; Okada and Hickok 2009; Isenberg et al. 2012; Venezia et al. 2016). We expected that language-selective subregions of Broca’s area and the PTL (identified in individual subjects) would exhibit increased activation for syntactically structured materials relative to unstructured word and nonword lists, consistent with previous findings. However, we hypothesized that production and perception would differentially recruit these regions: language-selective subregions of Broca’s area would be preferentially recruited by production, and that language-selective subregions of the PTL would be more equally driven by production and perception.

## Materials & Methods

### Subjects

We recruited 20 healthy, right-handed, native speakers of English with no history of neurological disfunction (age 18-32, average 21.9). Subjects were paid $25 an hour for two hours of participation, for a total of $50 in total compensation. All subjects gave informed consent to participate, and all procedures were approved by the Institutional Review Board of the University of South Carolina.

### Stimuli

The experiment was comprised of a 3 × 3 design: three different tasks (perceive+rest, perceive+rehearse, continuous perceive) by three levels of Content (PHONOLOGICAL, SYNTACTIC, LEXICAL). As we discuss in more detail later, for statistical comparisons we performed a used the contrast of perceive+rehease > perceive+rest to define the effect of PRODUCTION, whereas continuous perceive vs. rest was used to define the effect of PERCEPTION.

#### Phonological stimuli

We created the PHONOLOGICAL materials by generating pseudowords for which no syntactic category was obvious. We created 16 total pseudowords, roughly distributed across different speech segments divided into two sets, with the constraint of syllable structure CV(C), a single onset consonant with an optional single code consonant. Set 1 (initial position): *perwoth*, *nansow*, *ninyo*, *denferr*, *bulbom*, *nillex*, *seenig*, *tringess*. Set 2 (second position): *lerris*, *foyrix*, *pobset*, *ganliff*, *demesh*, *garlay*, *susset*, *furgle*. Each pseudoword from the first set was paired once with all of the pseudowords from the second set, producing 64 unique two-pseudoword sequences (e.g. *perwoth lerris*), with the first word presented on top of the second word on the screen during the experiment. We then repeated these 64 sequences for use across the entire experiment, copying the set of 64 once and then randomly selecting an additional subset of 22/64 sequences for a total of 150 sequences. We then distributed these 150 sequences randomly to create 30 trials each for the three task conditions, using one stimulus for each perceive+rest trial, one stimulus for each perceive+rehearse trial, and three stimuli for each continuous perceive trial.

#### Syntactic stimuli

We created the SYNTACTIC materials by adapting the phonologically plausible pseudoword nouns created by Matchin et al. (2017) using the Wuggy software (Keuleers & Brysbaert, 2010). This study created phonologically plausible pseudowords (nouns) preceded by real determiners to create phrases that preserved syntactic structure but greatly reducing conceptual content in order to investigate the neural bases of syntactic processing. They were designed to match the phonological plausibility of real nouns used within that study. We selected monosyllabic pseudoword nouns such that each jabberwocky phrase contained two syllables to match the phonological condition. The monosyllabic pseudowords (C)CVC(C), that is, minimum CVC with an optional additional onset OR coda consonant. i.e., the three possible syllables were CVC, CCVC, and CVCC. The final set was: *bleff*, *woon*, *pand*, *delk*, *sheeve*, *glit*, *lart*, *clope*. We then combined these pseudowords with a set of eight determiners to create 64 unique phrases: the articles *a* and *the*, possessive pronouns *his* and *their*, demonstratives *this* and *those*, and the quantifiers *each* and *few*. In order to ensure variability in syntactic number features across stimuli, *a*, *their*, *this*, and *each* were combined with a singular noun (e.g. *a bleff*) while *the*, *his*, *those*, and *few* were combined with a plural noun (e.g. *the pands*). In order to create stimuli that matched the phonological condition in number of syllables, we combined two phrases together to form each individual stimulus, randomly assigned with the constraint that the two phrases not overlap in either the pseudoword or determiner and balanced to have equal numbers of each determiner in both the first and second phrases. This resulted in 64 two-phrase sequences (e.g. *these clopes this pand*), with the first phrase presented on top of the second phrase on the screen during the experiment. As with the PHONOLOGICAL stimuli, we duplicated this set of 64 and added 22/64 randomly selected phrases to create a total of 150 sequences. We then distributed these 150 sequences randomly to create 30 trials each for the three task conditions, using one stimulus for each perceive+rest trial, one stimulus for each perceive+rehearse trial, and three stimuli for each continuous perceive trial.

#### Lexical stimuli

We created the LEXICAL materials by combining two semantically unrelated nouns each consisting of two syllables. From a set of 16 nouns, we divded them into two sets. Set 1 (initial position): *hermit*, *ninja*, *pirate*, *poet*, *sheriff*, *mutant*, *glutton*, *hostage*. Set 2 (second position): *dogma*, *vodka*, *garbage*, *pistol*, *organ*, *fortress*, *scandal*, *robot*. Each pseudoword from the first set was paired once with all of the pseudowords from the second set, producing 64 unique two-word sequences (e.g. *hermit dogma*), with the first word presented on top of the second word on the screen during the experiment. We then repeated these 64 sequences for use across the entire experiment, copying the set of 64 once and then randomly selecting an additional subset of 22/64 sequences for a total of 150 sequences. We then distributed these 150 sequences randomly to create 30 trials each for the three task conditions, using one stimulus for each perceive+rest trial, one stimulus for each perceive+rehearse trial, and three stimuli for each continuous perceive trial.

### Procedure

The experiment consisted of two phases: a training phase during which subjects performed the task outside of the scanner with overt speech production, and the testing phase in which subjects performed the task covertly inside of the scanner. Stimuli were presented using PsychToolBox (Brainard, 1997; Pelli, 1997; Kleiner et al., 2007). Task presentation was identical for the training and testing phases; the only difference was whether the subject articulated overtly (training phase) or imagined speaking (testing phase). All trials involved the presentation of a cue for 1s: the word Read (in white font), cueing the subject to comprehend the stimulus but not articulate, used for both the perceive+rest and continuous perceive conditions, or the word Repeat (in green font), used for the perceive+repeat condition, cueing the subject to comprehend the stimulus and then repeat it three times during the delay phase. After the cue, a fixation cross was presented for 1s in the same color font as the cue. Following this, a written speech stimulus was presented for 2s. In the continuous perceive condition, two additional speech stimuli were presented for 2s each, followed by a white fixation cross for 2s before the next trial. In the perceive+rest condition, a white fixation cross was presented for 6s. In the perceive+repeat condition, the screen was blank for 4s during which the subject was trained to repeat the speech stimulus three times, followed by a white fixation cross for 2s before the next trial. A schematic of stimulus presentation is shown in Figure 1.

**Figure 1.**
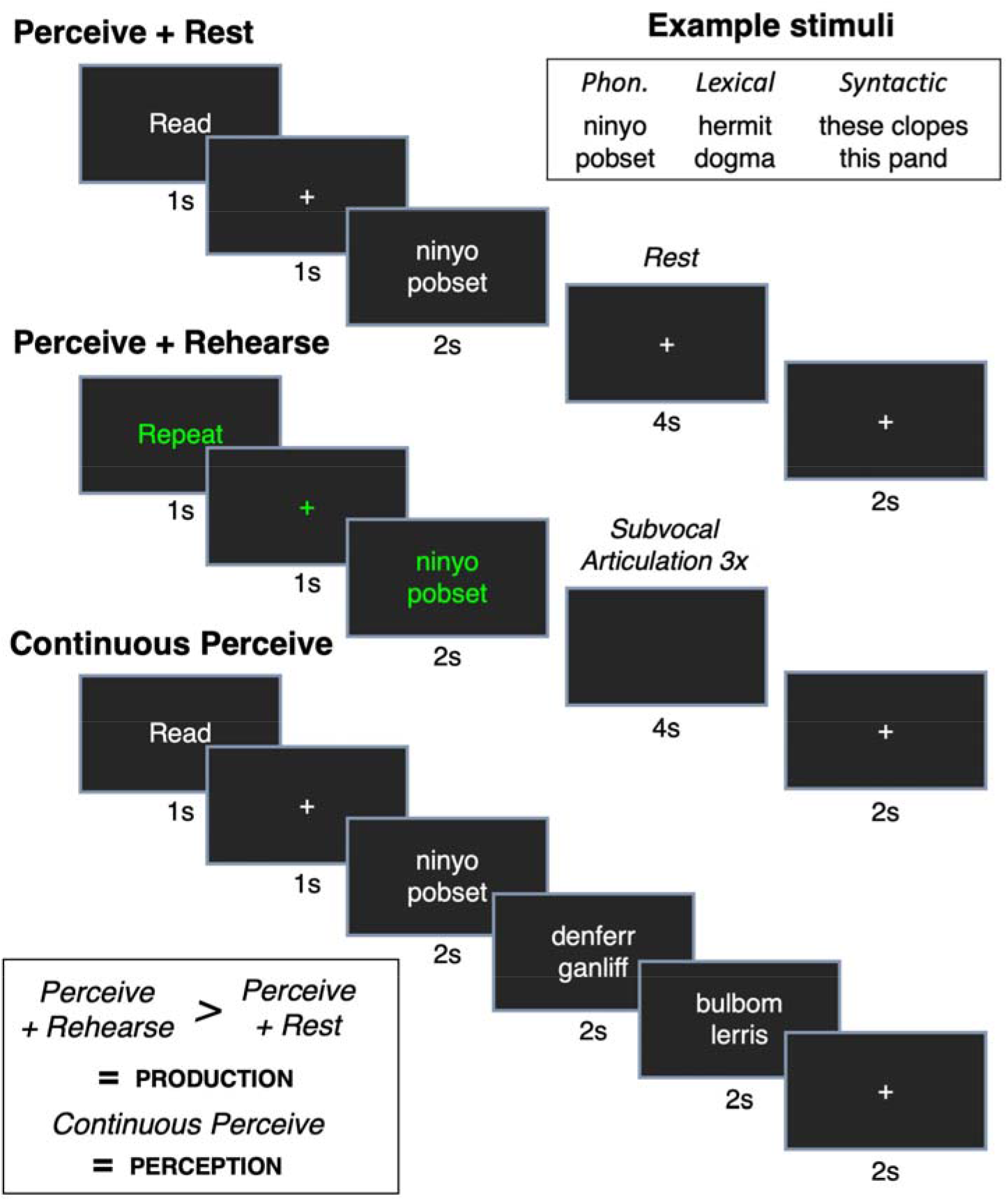
A schematic of stimulus design and presentation. See text for details. Phon. = phonological.

#### Training phase

During the training phase, the experiment was first explained and modeled to the subject by the experimenter. Subjects were told that when the word “repeat” appeared in green font (the perceive+rehearse condition) to produce the presented speech stimulus three times during the delay period between fixation crosses, during which no nothing appeared on the screen. For the perceive+rest and continuous perceive conditions, the subjects were told that when the word “read” appeared in white font to read and comprehend the speech stimuli presented on screen but not to produce anything. They were instructed to perform the task out loud during the training phase, but that they would only imagine speaking during the testing phase inside the scanner.

Following this, subjects performed the task in two short runs consisting of 24 trials each: six perceive+rest, six perceive+repeat, six continuous perceive, and six rest trials consisting of fixation only. Practice trials were randomly selected from the set of created stimuli and presented in random order, such that all conditions were balanced (that is, nine experimental conditions with two trials per condition). Random order was manually rearranged so that at least two non-rest trials intervened between rest trials, and runs always ended with a rest trial. The six overt perceive+rehearse trials per run, with three overt utterances per trial, were recorded for later analysis (for two subjects, auditory recordings were unavailable due to equipment issues, therefore only 18 subjects’ data were analyzed).

#### Testing phase

The experiment was divided into 9 runs of 40 trials each: ten perceive+rest, ten perceive+repeat, ten continuous perceive, and ten rest trials. The nine conditions (PHONOLOGICAL perceive+rest, PHONOLOGICAL perceive+rehearse, PHONOLOGICAL continuous perceive, LEXICAL perceive+rest, LEXICAL perceive+rehearse, LEXICAL continuous perceive, SYNTACTIC perceive+rest, SYNTACTIC perceive+rehearse, SYNTACTIC continuous perceive) were maximally balanced across runs. For example, in one run, for the ten perceive+rest trials, three were PHONOLOGICAL, three SYNTACTIC, and four LEXICAL. As in the practice run, at least two non-rest trials intervened between rest trials, and runs always ended on rest trials (to allow the BOLD response to return to baseline at the end of the run).

#### fMRI data collection and analysis

Brain data were obtained in a Siemens PRISMA 3T scanner (Siemens Medical Systems) using a 20-channel head coil. After the subjects were installed in the scanner, preliminary scans were obtained in order to localize the subject’s brain and adjust shim coils for magnetic field homogeneity. The subject was reminded not to produce any speech out loud but only to subvocally rehearse in the perceive+repeat trials. Following this, the subject performed four experimental runs, followed by a high resolution T1 anatomical scan, followed by the last five runs. Each run lasted approximately six minutes, and some subjects occasionally took a one minute break in-between runs. Following the last run, the subject was removed from the scanner, debriefed, and paid for their participation.

The high-resolution T1-weighted anatomical image was collected in the axial plane (voxel dimension: 1 mm isotropic) using an MP-RAGE sequence (256 × 256 matrix size, 9 degree flip angle). A total of 2880 T2*-weighted EPI volumes were collected over 9 runs of 320 volumes apiece. Each volume consisted of 68 slices in ascending, interleaved order without gap (TR = 1260 ms, TE = 32 ms, flip angle = 45°, in-plane resolution = 2.5 mm × 2.5 mm, slice thickness = 2.5 mm with no gap). The first four volumes of each run (dummy volumes) were discarded automatically by the scanner to control for T1 saturation effects. Data were reconstructed using MRIcroGL (https://www.nitrc.org/projects/mricrogl). Slice-timing correction, motion correction, warping to MNI space, spatial smoothing, and conversion to percent signal change values were performed using AFNI software (Cox 1996) http://afni.nimh.nih.gov/afni). Motion correction was achieved by using a 6-parameter rigid-body transformation, with each functional volume in each run first aligned to a single volume in that run. Functional volumes were aligned to the anatomical image, aligned to MNI space, and resampled to 3 mm isotropic. Functional images were spatially smoothed using a Gaussian kernel of 6 mm FWHM.

First-level (individual subject) analysis was performed for each subject using AFNI’s 3dDeconvolve function. The regression equation identified parameter estimates that best explained variability in the data, using a canonical hemodynamic response function convolved with the timing of stimulus presentation for each condition. We included a regressor for each of the nine conditions (PHONOLOGICAL perceive+rest, PHONOLOGICAL perceive+rehearse, PHONOLOGICAL continuous perceive, LEXICAL perceive+rest, LEXICAL perceive+rehearse, LEXICAL continuous perceive, SYNTACTIC perceive+rest, SYNTACTIC perceive+rehearse, SYNTACTIC continuous perceive), modeling the duration between the onset of the speech stimulus until the final fixation cross (6s). We added a 2s regressor for the cues (Read vs. Repeat) that preceded each speech stimulus. Finally, we included the six motion parameters as regressors of no interest. We then performed first-level contrasts to identify the effect of PRODUCTION (perceive+rehearse > perceive+rest) for each level of Content (PHONOLOGICAL, LEXICAL, SYNTACTIC) within each subject. The effect of PERCEPTION for each level of Content was defined as the continuous perceive condition vs. rest.

We defined subject-specific functional ROIs within broader anatomical search spaces (Fedorenko et al., 2010; Rogalsky et al., 2015; Matchin et al., 2019). We first defined two localizer contrasts, orthogonal to our effects of interest, to identify language-selective subregions in individual subjects: all SYNTACTIC conditions (perceive+rest, perceive+rehearse, continuous perceive) compared to all LEXICAL conditions, [SYNTACTIC > LEXICAL], and all SYNTACTIC conditions compared to all PHONOLOGICAL conditions, [SYNTACTIC > PHONOLOGICAL]. We created anatomical search spaces by combining ROIs within the Johns Hopkins University atlas: Broca’s area (inferior frontal gyrus, pars triangularis and pars opercularis) and the PTL (posterior superior temporal gyrus and middle temporal gyrus). We then intersected the subject-specific contrast maps thresholded at p < 0.005 with the anatomical search spaces to result in four individual ROIs for each subject: Broca’s area [SYNTACTIC > LEXICAL], Broca’s area [SYNTACTIC > PHONOLOGICAL], PTL [SYNTACTIC > LEXICAL], and PTL [SYNTACTIC > PHONOLOGICAL].

We then analyzed orthogonal functional dissociations within these regions at the group level, averaging the t-statistic for each condition across the voxels included within the subject-specific ROIs. We performed six separate 2×2 ANOVAs, separately for each linguistic level of Content (PHONOLOGICAL, LEXICAL, and SYNTACTIC) and each Region defined by the localizer contrasts ([SYNTACTIC > LEXICAL], [SYNTACTIC > PHONOLOGICAL]). We analyzed the main effects of Task (PERCEPTION vs. PRODUCTION), Region (Broca’s area, the PTL), and their interaction.

Our whole brain analyses were used to show the broader patterns of activation associated with our experimental manipulations. We created overlap maps to identify regions that showed increased activation for [SYNTACTIC > LEXICAL] and [SYNTACTIC > PHONOLOGICAL], separately for PRODUCTION and PERCEPTION, using a voxel-wise threshold of p < 0.001, cluster size 40 voxels, which resulted in reported analyses passing an FDR correction for multiple comparisons at q < 0.05. In Supplementary Materials, we show whole-brain activations for all of the 6 individual effects across the 2 tasks (PERCEPTION, PRODUCTION) x 3 linguistics levels (PHONOLOGICAL, LEXICAL, SYNTACTIC) design using these same statistical thresholds and FDR correction.

## Results

### Behavioral

Subjects performed well overall on attempting and accurately producing the presented speech stimuli during the training period prior to scanning (Figure 2): 94% of attempted productions were accurate in the PHONOLOGICAL condition, 88% of attempted productions were accurate in the SYNTACTIC condition, and 98% of attempted productions were accurate in the LEXICAL condition. Paired samples t-tests revealed significantly better accuracy for PHONOLOGICAL read+repeat relative to SYNTACTIC read+repeat, *t*(1,17) = 2.787, *p* = 0.013, significantly better accuracy for read+repeat relative to SYNTACTIC read+repeat, *t*(1,17) = 4.136, *p* < 0.001, and significantly better accuracy for LEXICAL read+repeat relative to PHONOLOGICAL read+repeat, *t*(1,17) = 4.526, *p* < 0.001.

**Figure 2.**
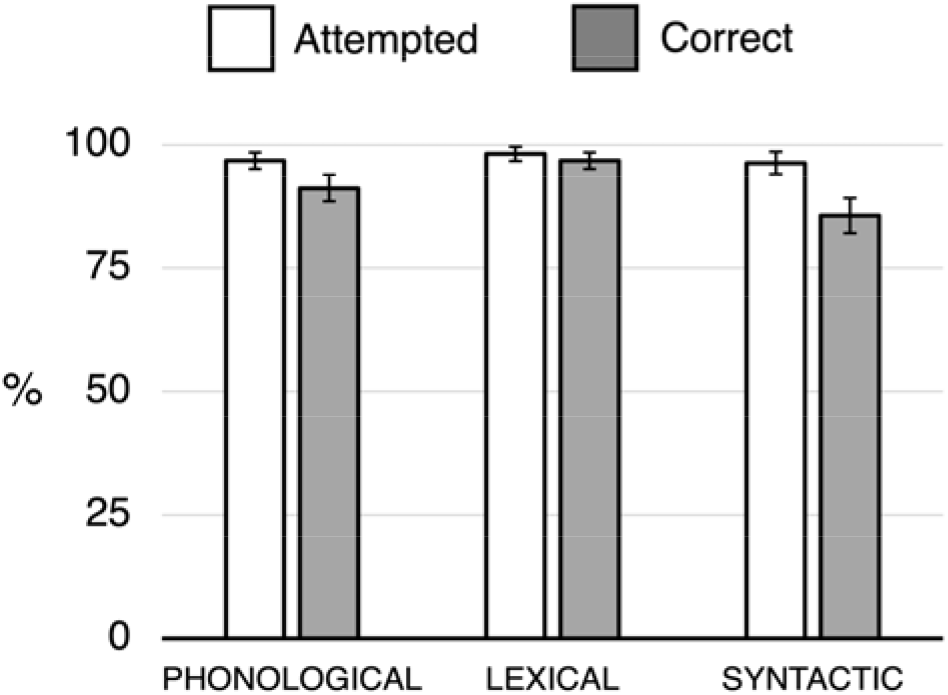
Behavioral data from the training phase. Data are shown as a percentage of the total possible number of utterances that were attempted and made correctly for each of the three levels of Content (PHONOLOGICAL, LEXICAL, SYNTACTIC) during the perceive+rehearse task. Statistical analyses were performed on the proportion of accurate/attempted utterances.

### fMRI - Whole brain analyses

At the whole brain level, no significant clusters emerged using for syntactic effects in PRODUCTION, the [SYNTACTIC > LEXICAL] and [SYNTACTIC > PHONOLOGICAL] contrasts. Significant clusters for syntactic effects in PERCEPTION are shown in Figure 3. The [SYNTACTIC > LEXICAL] contrast revealed activation in a broad set of regions, including left anterior precentral gyrus extending into the posterior inferior frontal gyrus; superior precentral gyrus; left pSTS/MTG; bilateral posterior ventral occipito-temporal cortex, bilateral dorsal occipital/parietal lobe, and bilateral calcarine sulcus (Figure 3, dark blue). The [SYNTACTIC > PHONOLOGICAL] contrast activated essentially a subset of these regions, including pSTS/MTG, bilateral calcarine sulcus, and dorsal occipital/parietal lobe (Figure 3, yellow). Overlap between these effects was observed in left pSTS/MTG, bilateral calcarine sulcus, bilateral thalamus, and bilateral dorsal occipital/parietal lobe (Figure 3, red). Center of mass coordinates for these effects are listed in Table 1.

**Figure 3.**
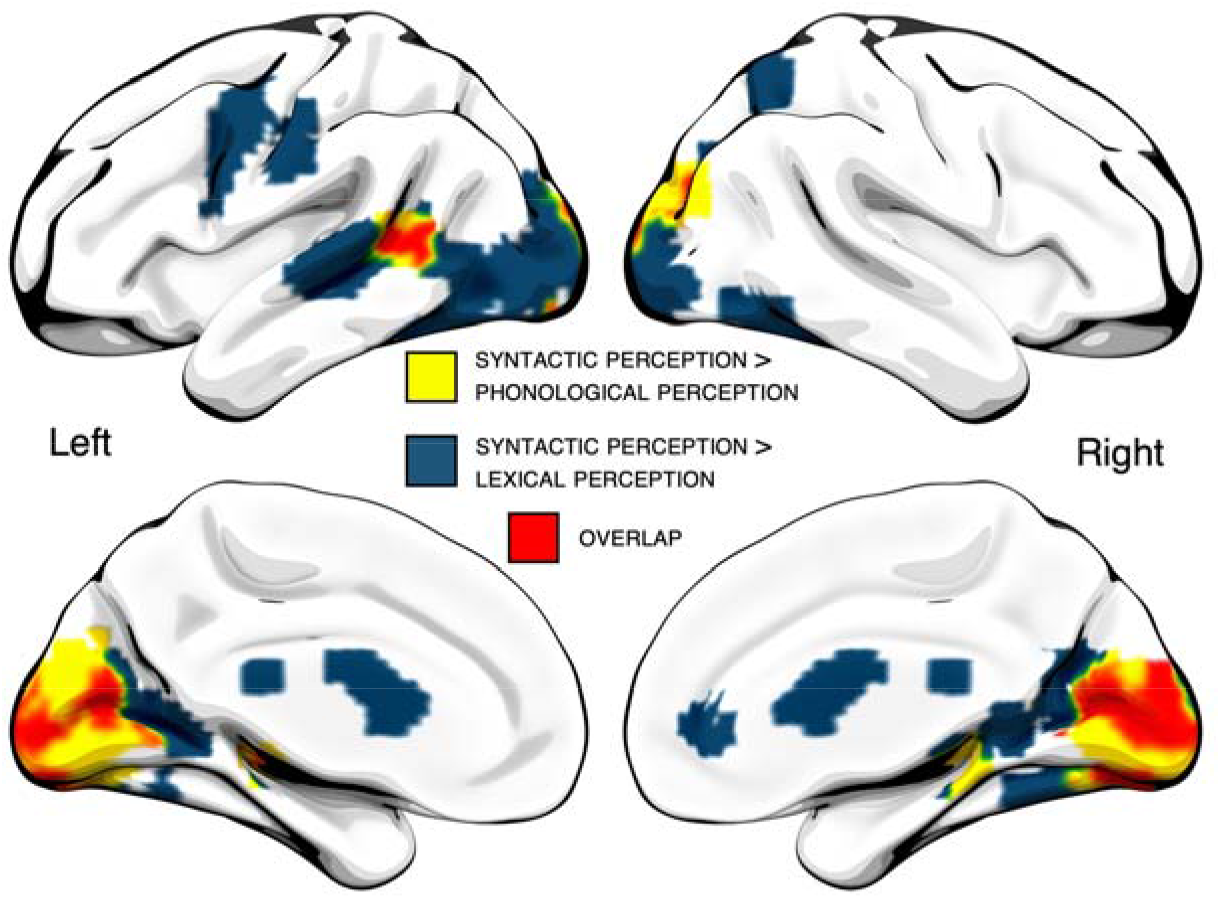
Conjunction (overlap) analysis of [SYNTACTIC > LEXICAL] and [SYNTACTIC > PHONOLOGICAL] in PERCEPTION at the whole brain level.

**Table 1.** Center of mass coordinates, reported in MNI space, for clusters of increased activation for [SYNTACTIC > PHONOLOGICAL] and [SYNTACTIC > LEXICAL] in PERCEPTION in the whole brain conjunction/overlap analysis.

| Region | Size | X | Y | Z |
| --- | --- | --- | --- | --- |

| SYNTACTIC PERCEPTION > PHONOLOGICAL PERCEPTION |  |  |  |  |
| --- | --- | --- | --- | --- |
| Bilateral calcarine sulcus | 61,884 mm<sup>3</sup> | 1 | -84 | 4 |
| Left pSTS/MTG | 4,293 mm<sup>3</sup> | -54 | -49 | 9 |
| Right thalamus | 3,375 mm<sup>3</sup> | 15 | -30 | -9 |
| Left thalamus | 1,566 mm<sup>3</sup> | -21 | -24 | -4 |
| SYNTACTIC PERCEPTION > LEXICAL PERCEPTION |  |  |  |  |
| Bilateral calcarine sulcus | 76,464 mm<sup>3</sup> | -7 | -75 | 1 |
| Bilateral basal ganglia | 16,929 mm<sup>3</sup> | -3 | -25 | 17 |
| Left precentral gyrus | 9,828 mm<sup>3</sup> | -49 | -3 | 41 |
| Right cerebellum | 2,052 mm<sup>3</sup> | 25 | -63 | -53 |
| Right superior parietal lobule | 1,566 mm<sup>3</sup> | 26 | -58 | 55 |
| Right thalamus | 1,404 mm<sup>3</sup> | 19 | -26 | -2 |
| Left thalamus | 1,404 mm<sup>3</sup> | -22 | -26 | -4 |
| Right anterior cingulate cortex | 1,107 mm<sup>3</sup> | 12 | 39 | 4 |
| *Overlap* |  |  |  |  |
| Bilateral calcarine sulcus | 7,722 mm<sup>3</sup> | 3 | -84 | 2 |
| Left pSTS/MTG | 2,538 mm<sup>3</sup> | -56 | -49 | 9 |
| Left thalamus | 1,080 mm<sup>3</sup> | -21 | -25 | -4 |
| Right thalamus | 621 mm<sup>3</sup> | 20 | -27 | -3 |

### fMRI - ROI analyses

The four subject-specific functional ROIs are shown in Figure 4. For the localizer contrast [SYNTACTIC > LEXICAL], maximum overlap in Broca’s area (11 subjects) occurred in the pars opercularis, MNI peak coordinates [−47 10 22], and maximum overlap in the PTL (13 subjects) occurred in the superior temporal sulcus, MNI peak coordinates [−53, −41, 7]. For the localizer contrast [SYNTACTIC > PHONOLOGICAL], one subject did not have significant voxels in Broca’s area, so we omitted this subject’s data in all analyses for this functional localizer. Maximum overlap in Broca’s area (7 subjects) occurred in 9 mostly noncontiguous voxels, with a general bias towards the pars triangularis (6 peak voxels) rather than pars opercularis (3 peak voxels), and maximum overlap in the PTL (14 subjects) occurred in the superior temporal sulcus, MNI peak coordinates [−58, −50, 8]. Overall, there was a high degree of correspondence between the two localizer contrasts in the PTL, but in Broca’s area there was a noticeable distinction between the [SYNTACTIC > LEXICAL] contrast (posterior) and the [SYNTACTIC > PHONOLOGICAL] contrast (anterior).

**Figure 4.**
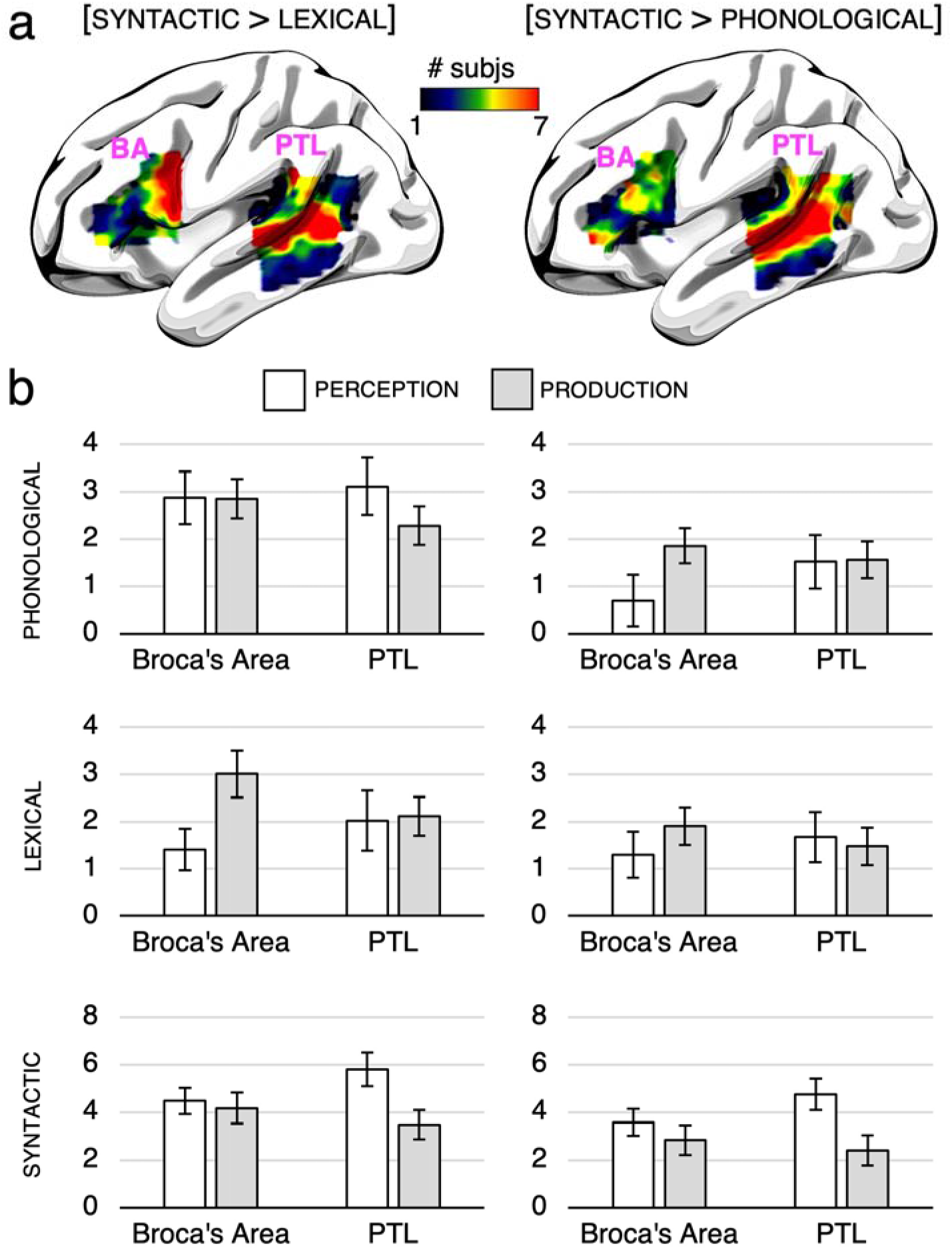
(a): language-selective ROIs defined in individual subjects. Color indicates the number of subjects with overlapping significant voxels in each ROI. (a, left): ROIs defined by the localizer contrast [SYNTACTIC > LEXICAL]. (a, right): ROIs defined by the localizer contrast [SYNTACTIC > PHONOLOGICAL]. (b): Bar charts showing the average t-value within each ROI for PERCEPTION and PRODUCTION at each linguistic level. Bar charts on the left correspond to the functional ROIs defined using the [SYNTACTIC > LEXICAL] contrast, and bar charts on the right correspond to the functional ROIs defined using the [SYNTACTIC > PHONOLOGICAL] contrast. Error bars indicate standard error of the mean. BA = Broca’s area, PTL = posterior temporal lobe.

The average t-statistic for each condition within each ROI is shown in Figure 4. For all ROIs and linguistic levels of Content, Broca’s area had higher activation for PRODUCTION relative to the PTL, and PTL had higher activation for PERCEPTION relative to Broca’s area. Table 2 contains the statistical results of the analyses performed within these ROIs. For the main effects of Task (SYNTACTIC PERCEPTION vs. SYNTACTIC PRODUCTION) and Region (Broca’s area vs. PTL), our analyses revealed no significant differences, indicating that there were no overall differences in activation between Broca’s area and the PTL or between SYNTACTIC PERCEPTION and SYNTACTIC PRODUCTION. There was a highly significant interaction between Task and Region for both sets of ROIs, p < 0.001. Thus there was robust evidence for a production-perception asymmetry between language-selective subregions of Broca’s area and PTL for syntactic processing. Follow-up pairwise comparisons revealed no significant differences between syntactic production between Broca’s area and the PTL, and marginally significant differences between syntactic perception between Broca’s area and the PTL (not surviving a Bonferroni correction for multiple comparisons). This suggests that these regions responded roughly equally for production, but that PTL showed enhanced activity for perception, consistent with the predictions of Matchin & Hickok (2020).

**Table 2.** Statistical results of the ANOVAs comparing syntactic production and syntactic perception between the functionally-defined, subject-specific ROIs in Broca’s area and the posterior temporal lobe (PTL). * indicates significance before multiple comparisons correction, and bolding indicates significance surviving a Bonferroni correction for multiple comparisons (four pairwise comparisons, p < 0.0125).

|  | [SYNTACTIC > LEXICAL] ROI |  | [SYNTACTIC > PHONOLOGICAL] ROI |  |
| --- | --- | --- | --- | --- |
| Region | F(1,19) = 0.524 | p = 0.478 | F(1,18) = 1.012 | p = 0.328 |
| Task | F(1,19) = 2.012 | p = 0.172 | F(1,18) = 3.674 | p = 0.071 |
| **Interaction** | **F(1,19) = 19.524** | **\*p < 0.001** | **F(1,18) = 15.945** | **\*p < 0.001** |
| Effect of Region for SYNTACTIC PRODUCTION | t(1,19) = 1.881 | p = 0.075 | t(1,18) = 0.501 | p = 0.622 |
| Effect of Region for SYNTACTIC PERCEPTION | t(1,19) = -2.287 | *p = 0.034 | t(1,18) = -2.173 | *p = 0.043 |

## Discussion

The lack of strong differentiation among regions with respect to syntactic effects in neuroimaging studies in the literature has led some to call into question claims of localization of syntactic processing to a single brain area (Fedorenko et al. unpublished preprint; Kaan and Swaab 2002; Blank et al. 2016). Particularly striking is the high degree of similarity of syntactic activation profiles between Broca’s area and the posterior temporal lobe (PTL) (see Matchin & Hickok, 2020 for a review). Consistent with these observations, we identified syntax-sensitive subregions of both of these areas in individual subjects. However, we identified a clear asymmetry with respect to syntactic demands in production and perception: activation in Broca’s area was driven more by the demands of production than PTL, and activation in the PTL was driven more by the demands of perception than Broca’s area. This suggests that these regions make distinct contributions to syntactic processing.

With respect to precise localization, our two individual subject localizer contrasts, [SYNTACTIC > LEXICAL] and [SYNTACTIC > PHONOLOGICAL], identified similar pSTS/MTG regions, both of which have been identified in previous studies of syntactic processing (e.g. Pallier et al., 2011; Matchin et al., 2017; 2019). At the group level, these contrasts overlapped in pSTS, suggesting a fairly robust role for this region in syntactic processing. However, our two localizer contrasts identified distinct subregions of Broca’s area: the [SYNTACTIC > LEXICAL] contrast highlighted pars opercularis, while the [SYNTACTIC > PHONOLOGICAL] contrast highlighted pars triangularis. Additionally, at the group level, only the [SYNTACTIC > LEXICAL] contrast found a significant effect in pars opercularis, with no significant effects for the [SYNTACTIC > PHONOLOGICAL] contrast. While some studies have identified primary activation foci for syntactic processing in the pars triangularis (Pallier et al. 2011; Matchin et al. 2017; Matchin, Liao, et al. 2019), other studies have identified primary foci in the pars opercularis (e.g. Goucha et al., 2015; Zaccarella et al., 2017). The Matchin & Hickok (2020) model suggests that the key subregion for morpho-syntactic processing in Broca’s area is the pars triangularis, based on the fact that phonological production demands (Buchsbaum et al., 2001; Hickok et al., 2003; Matchin et al., 2014) and phonological working memory demands in comprehension (Matchin et al., 2017; 2019) appear to drive activity in pars opercularis. We suggest that future research investigation the relationship between phonological processing and syntactic effects in pars opercularis be more thoroughly investigated.

One objection to the conclusion that syntax-sensitive regions of Broca’s area are differentially driven by production/comprehension demands relative to the PTL is the hypothesis that this region contains distinct subregions, some of which are sensitive to higher-level syntax, some of which are sensitive to production, and that they are interdigitated, making it difficult to disentangle them. However, we identified our ROIs in individual subjects using language-selective localizer contrasts. Thus the interdigitated explanation is unlikely (although not impossible to rule out, if the interdigitation is finer than our voxel resolution), suggesting instead that language-selective regions of Broca’s area reflect a distinct linguistic computation from that of the PTL.

While Broca’s area has been implicated in production since the 1800s (Broca 1961), this role is not restricted to articulatory and/or phonological demands, at least in the pars triangularis. Recent electrocorticography studies of speech production have revealed that Broca’s area is not active during speech articulation (Flinker et al. 2015), and is implicated in higher-level morphological processes (Moro et al. 2001; Sahin et al. 2009). Matchin & Hickok (2020) recently proposed that the pars triangularis area underlies a morpho-syntactic sequencing function, tied to the demands of production, whereas the posterior superior temporal sulcus/middle temporal gyrus (pSTS/MTG) is critically involved in hierarchical lexical-syntactic structuring, supporting both comprehension and production. The functional asymmetry that we observed in the present study is consistent with this proposal.

If the contribution of Broca’s area to syntactic processing is mostly driven by the demands of production, why did we observe a significant activation for syntactic processing in perception in this region? It is unlikely that this activation reflects working memory demands (cf. (Rogalsky and Hickok 2011), as our stimuli involved maximally simple sequences of two-word phrases. We suggest, in agreement with other authors, that activation in Broca’s during perception in our task may reflect the prediction of upcoming material (Bonhage et al., 2015; Matchin et al., 2017; Matchin, 2018; Rimmele et al., 2018). A role in top-down predictions is supported by the fact that lesions to IFG impair the rapid processing of syntactic violations (Jakuszeit et al. 2013), and when sentence presentation is slowed, patients with IFG lesions show improved comprehension (Love et al. 2008). Thus some activation for syntactic perception is expected in this region, albeit asymmetrically with respect to the PTL, which appears to underlie core computations necessary for successful comprehension.

While we expected the PTL to be engaged roughly equally by both production and perception, the PTL ROIs appeared to show increased activity for perception relative to production. This was not predicted, particularly given a recent study which showed that paragrammatism in speech production, that is the misuse of syntactic structure and functional elements, is associated primarily with posterior temporal-parietal damage (Matchin et al., 2020). However, it may be that the articulatory rehearsal paradigm we used in the present study did not force subjects to always reproduce morpho-syntactic representations as opposed to purely phonological ones. In the rehearsal condition, some subjects may have converted the syntactic representation to a purely phonological one and rehearsed a phonological, rather than morpho-syntactic, sequence. By contrast, the demands of natural sentence production require generating variable morpho-syntactic sequences. Future research using this paradigm should ensure that speech sequences cannot be rehearsed purely in a phonological code, but rather a task should be implemented that requires subjects to re-code the morpho-syntactic structure of the utterance. We would predict that under such circumstances, activation in PTL will be equivalent for perception and production, and there will be increased activation for production relative to perception in Broca’s area.

Finally, although our materials were matched for number of syllables, our whole brain analyses revealed effects in the occipital lobe that were likely due to differences in the visual display among the conditions. Future research using a similar experimental design should explore other modalities of presentation, such as auditory speech and sign language. Previous studies using these disparate modalities have illustrated similar effects with tight overlap in the pSTS/MTG and Broca’s area (MacSweeney 2002; MacSweeney et al. 2006; Spitsyna et al. 2006; Jobard et al. 2007; Lindenberg and Scheef 2007; Pa et al. 2008; Berl et al. 2010; Leonard et al. 2012; Vagharchakian et al. 2012; Regev et al. 2013; Wilson et al. 2018). Therefore we would expect the same production-perception asymmetry across modalities in these regions.

### Conclusions

Our results point towards a possible resolution of a conflict between the functional neuroimaging literature on syntax, which has not identified robust differences between Broca’s area and the PTL in syntactic comprehension, and the lesion-symptom mapping literature, which has identified multiple dissociations with respect to damage to these regions. Namely, that activation in language-selective regions of the PTL are driven by the demands of hierarchical structure-building necessary to comprehend the meaning of a sentence, whereas language-selective regions of Broca’s area are driven by the demands of production, such as converting a structure into a linear string of morphemes. Future neuroimaging studies should seek to provide more direct evidence of a specific functional role for Broca’s area in production-related processes.

## Funding

This research was supported by startup funds at the University of South Carolina awarded to William Matchin, and a Magellan Scholar Award. from the University of South Carolina awarded to Emily Wood and William Matchin.

## Notes

### Competing Interest Statement

The authors have declared no competing interest.

