## Supplementary Materials for "Syntax-sensitive regions of Broca’s area and the posterior temporal lobe are differentially recruited by production and perception"

Individual activation maps associated with the effect of production (perceive+rehearse > perceive+rest) for each linguistic level of Content (phonological, lexical, syntactic) are shown in Supplementary Figure 1, and individual activation maps associated with the effect of perception (continuous perceive > rest) for each linguistic level of Content (phonological, lexical, syntactic) are shown in Supplementary Figure 2. Activations were similar among the levels of content. For the effects of production, all linguistic levels of Content revealed activation in posterior IFG (pars opercularis), precentral gyrus, inferior parietal lobe, posterior STS/MTG, medial occipital cortex, and supplementary motor area (SMA). Activations were bilateral but stronger in the left hemisphere, particularly in frontal and parietal cortex. For the effects of perception, all levels of content revealed activation in left inferior frontal sulcus, left dorsal precentral gyrus, left SMA, and bilateral ventro-lateral occipital-temporal cortex. Additionally, syntactic perception produced significant clusters in left pSTS and right precentral gyrus, and lexical perception produced significant clusters in bilateral basal ganglia and small activations in right precentral gyrus.

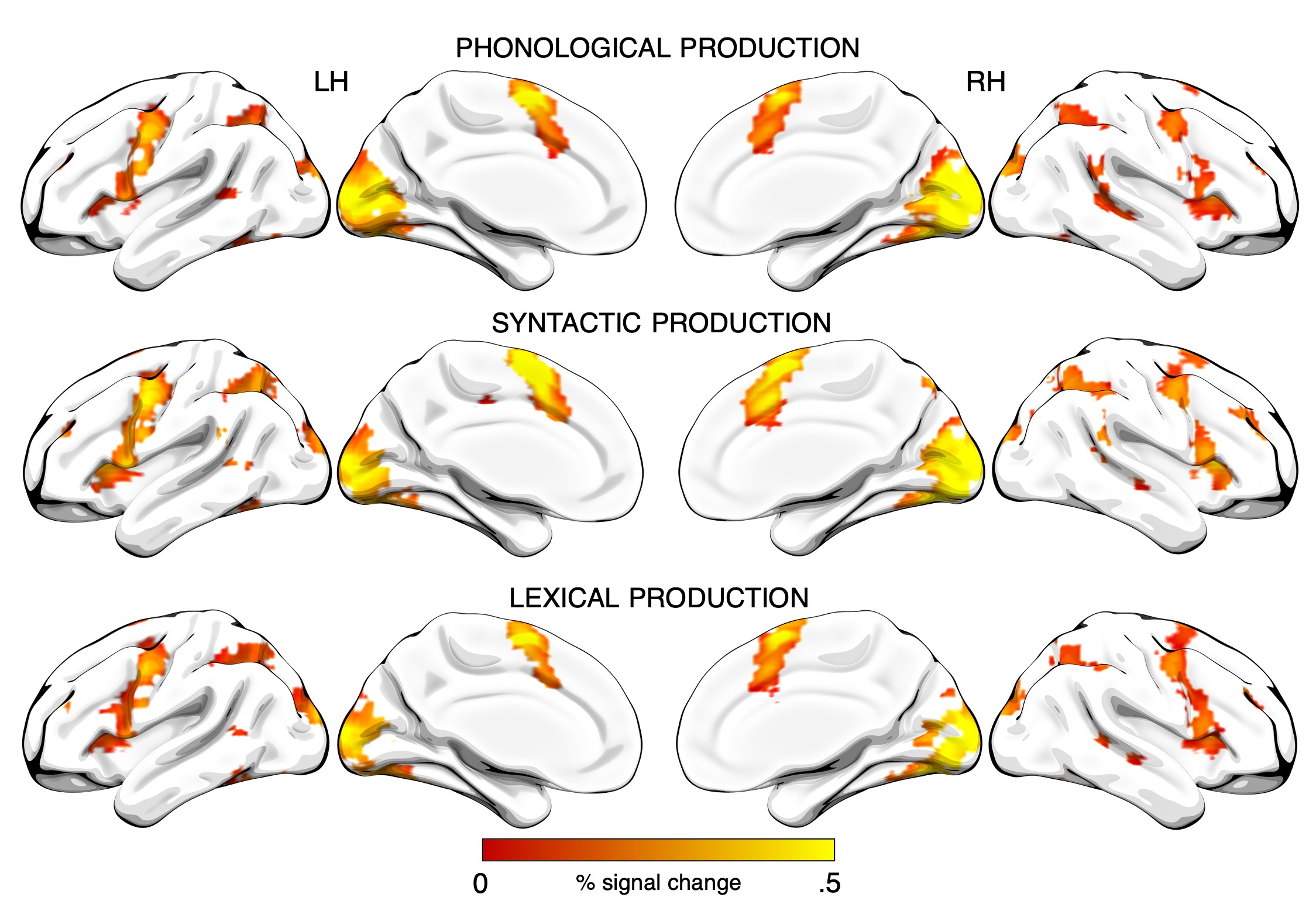

Figure 1. Significant clusters for the effect of production at each linguistic level of Content (phonological, lexical, syntactic) shown on an inflated brain in MNI space.

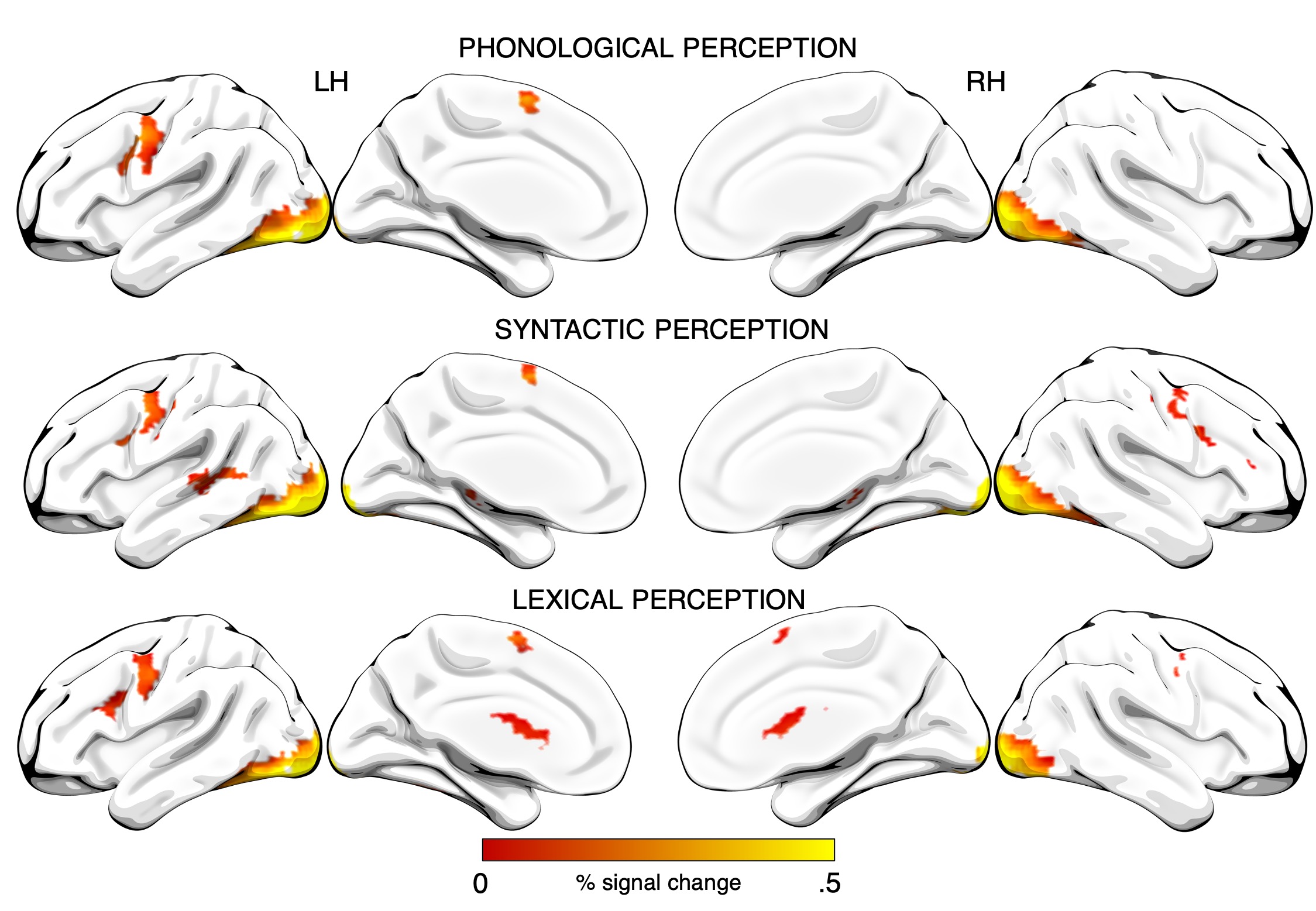

Figure 2. Significant clusters for the effect of perception at each linguistic level of Content (phonological, lexical, syntactic) shown on an inflated brain.

**Full list of materials**

Below is a list of the 64 unique materials for each linguistic level of Content that were used to generate the larger body of stimuli in the experiment.

*Phonological*

seenig	pobset

nillex	lerris

ninyo	lerris

denferr	pobset

seenig	furgle

bulbom	garlay

tringess	ganliff

denferr	garlay

denferr	lerris

nansow	lerris

seenig	susset

ninyo	demesh

denferr	ganliff

nansow	pobset

nansow	furgle

perwoth	ganliff

ninyo	furgle

nillex	susset

tringess	foyrix

seenig	lerris

seenig	ganliff

nansow	demesh

nillex	garlay

perwoth	susset

bulbom	foyrix

bulbom	demesh

perwoth	furgle

tringess	furgle

bulbom	lerris

perwoth	pobset

nillex	furgle

ninyo	foyrix

seenig	foyrix

tringess	garlay

denferr	susset

perwoth	demesh

nillex	foyrix

tringess	lerris

bulbom	ganliff

nansow	ganliff

perwoth	foyrix

perwoth	garlay

seenig	garlay

seenig	demesh

denferr	foyrix

ninyo	pobset

ninyo	susset

bulbom	furgle

nansow	garlay

nillex	ganliff

nillex	demesh

denferr	furgle

bulbom	susset

denferr	demesh

bulbom	pobset

tringess	demesh

perwoth	lerris

ninyo	ganliff

nillex	pobset

ninyo	garlay

nansow	foyrix

tringess	susset

tringess	pobset

nansow	susset

*Lexical*

pirate	garbage

mutant	pistol

mutant	garbage

hermit	pistol

glutton	dogma

sheriff	organ

glutton	vodka

sheriff	robot

glutton	robot

pirate	scandal

poet	scandal

sheriff	pistol

mutant	organ

poet	organ

mutant	scandal

hermit	robot

sheriff	dogma

ninja	organ

hostage	pistol

mutant	robot

ninja	fortress

hermit	organ

pirate	robot

hostage	vodka

mutant	vodka

ninja	vodka

hostage	robot

ninja	garbage

pirate	organ

mutant	dogma

pirate	dogma

hermit	vodka

ninja	scandal

poet	robot

pirate	fortress

sheriff	vodka

mutant	fortress

glutton	pistol

hostage	fortress

sheriff	garbage

hermit	fortress

hermit	dogma

hermit	scandal

poet	pistol

sheriff	fortress

sheriff	scandal

hostage	garbage

hostage	organ

pirate	pistol

glutton	organ

hostage	scandal

glutton	scandal

poet	garbage

pirate	vodka

ninja	robot

glutton	garbage

ninja	dogma

hermit	garbage

poet	dogma

poet	vodka

ninja	pistol

glutton	fortress

poet	fortress

hostage	dogma

*Syntactic*

few	sheeves	those	bleffs

his	larts	the	sheeves

each	pand	a	clope

the	delks	his	pands

his	glits	few	pands

each	woon	those	sheeves

their	woon	each	glit

a	glit	the	bleffs

his	delks	their	bleff

the	pands	his	bleffs

a	sheeve	the	woons

this	woon	those	pands

his	pands	each	woon

this	lart	their	glit

few	woons	those	clopes

this	sheeve	a	woon

few	glits	the	larts

few	pands	those	larts

a	pand	those	delks

few	clopes	those	glits

this	clope	his	sheeves

their	delk	his	glits

each	glit	those	woons

his	sheeves	this	glit

his	bleffs	the	clopes

those	clopes	few	larts

their	bleff	the	delks

their	clope	this	pand

the	clopes	their	lart

this	bleff	each	delk

a	clope	this	bleff

a	lart	this	clope

a	woon	each	pand

the	woons	each	lart

those	delks	their	sheeve

their	pand	his	delks

each	clope	their	delk

a	delk	his	clopes

each	delk	their	pand

each	sheeve	a	pand

this	pand	their	woon

those	sheeves	his	woons

those	larts	each	clope

his	clopes	this	woon

the	bleffs	this	lart

each	bleff	a	lart

a	bleff	this	delk

those	bleffs	the	glits

their	glit	few	bleffs

the	glits	each	bleff

those	pands	few	delks

his	woons	each	sheeve

the	larts	a	delk

their	sheeve	a	bleff

few	bleffs	a	sheeve

those	glits	few	sheeves

few	larts	this	sheeve

this	delk	their	clope

this	glit	the	pands

their	lart	a	glit

the	sheeves	few	woons

few	delks	his	larts

each	lart	few	glits

those	woons	few	clopes
